# APC/C ^Cdh1^-mediated degradation of Cdh1 during meiosis in *S. cerevisiae*

**DOI:** 10.1101/358655

**Authors:** Denis Ostapenko, Mark J. Solomon

## Abstract

The Anaphase-Promoting Complex/Cyclosome (APC/C) is a ubiquitin ligase that promotes the ubiquitination and subsequent degradation of numerous cell cycle regulators during mitosis and in G1. Proteins are recruited to the APC/C by activator proteins such as Cdh1. During the cell cycle, Cdh1 is subject to precise regulation so that substrates are not degraded prematurely. We have explored the regulation of Cdh1 during the developmental transition into meiosis and sporulation in the budding yeast *S. cerevisiae*. Transition to sporulation medium triggers the degradation of Cdh1. Degradation requires that cells be of the a/a mating type and be starved for glucose, but they do not actually need to enter into the meiotic program. Degradation requires an intact SNF1 protein kinase complex (orthologous to the mammalian AMPK nutritional sensor), which is activated by the absence of glucose. Cdh1 degradation is mediated by the APC/C itself in a ‘*trans*’ mechanism in which one molecule of Cdh1 recruits a second molecule of Cdh1 to the APC/C for ubiquitination. However, Cdh1-Cdh1 recognition does not depend on the degradation motifs or binding sites involved in the recognition of typical APC/C substrates. We hypothesize that Cdh1 degradation is necessary for the preservation of cell cycle regulators and chromosome cohesion proteins between the reductional and equational meiotic divisions, which occur without the intervening Gap or S phases found in mitotic cell cycles.

## INTRODUCTION

Protein ubiquitination regulates the stability of numerous proteins controlling cell cycle progression and differentiation. During cell growth, protein ubiquitination is carried out by the coordinated action of an E1 (ubiquitin-activating enzyme), multiple E2s (ubiquitin-conjugating enzymes), and two major E3 complexes (ubiquitin ligases): Skp1-Cullin-F-box protein (SCF) and the anaphase-promoting complex or cyclosome (APC/C) (Cardozo and Pagano, 2004; Peters, 2006; Thornton and Toczyski, 2006; Barford, 2011; Pines, 2011; Primorac and Musacchio, 2013). The resulting covalent attachment of poly-ubiquitin chains target modified proteins for degradation by the 26S proteasome. The evolutionarily conserved APC/C is an essential RING-type ubiquitin ligase composed of a large catalytic core and a substrate-binding activator protein, Cdc20 and Cdh1 in vegetative cells and Ama1 in meiotic cells (Visintin *et al*., 1997; Barford, 2011; Primorac and Musacchio, 2013). The APC/C activators both bring protein substrates to the vicinity of the catalytic core and modulate APC/C enzymatic activity. Both Cdc20 and Cdh1 recognize substrates through direct binding to a degradation motif (degron), typically a Destruction box (D-box, RxxLxxxxN), a KEN motif, an ABBA motif, and their derivatives (Glotzer *et al*., 1991; Pfleger *et al*., 2001; Di Fiore *et al*., 2015). Each of these degradation motifs interacts with a corresponding receptor within the activator’s WD-40 domain, and mutations within these degradation motifs or their receptors prevent substrate recruitment and ubiquitination (He *et al*., 2013; Davey and Morgan, 2016; Qin *et al*., 2016). Following substrate binding, Cdc20 and Cdh1 associate with the Apc1, Cdc23 and Cdc27 APC/C subunits through two evolutionarily conserved motifs, the C-box within the protein and the invariant Ile-Arg (IR) residues at the C-terminus, with both contacts being essential for the efficient docking of activators and substrate ubiquitination (Brown *et al*., 2015; Chang *et al*., 2015).

To prevent unscheduled substrate degradation, the activities of Cdh1 and Cdc20 are tightly regulated by numerous post-translational modifications, association with inhibitory factors, and protein stability. During G1 phase, Cdc20 is degraded in an APC/C^Cdh1^-dependent manner and is also inhibited by the spindle assembly checkpoint, which ensures that all chromosomes are properly attached to the mitotic spindle before the onset of mitosis (Prinz *et al*., 1998; Shirayama *et al*., 1998; Foe *et al*., 2011; Foster and Morgan, 2012). Cdh1 is inhibited by phosphorylation by cyclin-dependent protein kinases (CDK), polo-like kinase Cdc5, and the meiosis-specific kinase Ime2 (Zachariae *et al*., 1998; Jaquenoud *et al*., 2002; Zhou *et al*., 2003; Holt *et al*., 2007; Crasta *et al*., 2008). CDK interacts with Cdh1 in a complex, mutually inhibitory manner: Cdh1 inhibits CDK by promoting the ubiquitination and degradation mitotic B-cyclins, whereas CDK-mediated phosphorylation near a nuclear localization signal (NLS) in Cdh1 and near its C-box cause its nuclear export and prevent its interaction with the core APC/C, respectively (Brandeis and Hunt, 1996; Irniger and Nasmyth, 1997; Schwab *et al*., 1997; Zachariae *et al*., 1998; Jaspersen *et al*., 1999; Hockner *et al*., 2016).

In a natural environment, yeast cells are often exposed to various stresses, including nutrient limitation. Under these conditions, diploid yeast cells can undergo a dramatic differentiation from vegetative growth to meiosis and sporulation, culminating in the production of four haploid spores capable of withstanding prolonged periods of starvation or dehydration (Neiman, 2011). The decision to initiate the meiotic program involves the integration of information from numerous signaling pathways, including those sensing limited carbon and nitrogen sources. Ultimately, APC/C-dependent ubiquitination of the Ume6 transcriptional repressor leads to transcriptional activation of *IME1* (*I*nducer of *ME*iosis) and expression of early meiotic genes (Kassir *et al*., 1988; Mallory *et al*., 2007). The ubiquitin ligase activity of APC/C is essential for all stages of meiosis but its regulation and substrate selectivity are substantially different from that in vegetative cells (Cooper and Strich, 2011). In contrast to mitosis, where mitotic cyclins are completely degraded in G1 phase by APC^Cdh1^, many APC/C substrates persist between meiosis I and meiosis II to prevent DNA re-replication and to maintain chromosome cohesion between the reductional division of meiosis I and the equatorial division of meiosis II. In addition, *S. cerevisiae* cells express a meiosis-specific APC/C activator, Ama1, a Cdh1 homolog that controls ubiquitination of key regulatory proteins (Cooper *et al*., 2000). Later in meiosis, Ama1-mediated ubiquitination promotes the degradation of Cdc20 and other regulatory proteins, including the polo-like kinase Cdc5, the mitotic cyclin Clb3, and the meiosis-specific transcription factor Ndt80 (Penkner *et al*., 2005; Tan *et al*., 2011; Okaz *et al*., 2012; Tan *et al*., 2013).

While the meiotic appearance of APC/C^Ama1^ and the resultant degradation of Cdc20 have been clear for some time, the fate of Cdh1 has been uncertain. We hypothesized that some mitotic substrates of APC/C^Cdh1^ may need to be protected from degradation during meiosis. In this manuscript, we report that Cdh1 levels decrease upon growth in sporulation-inducing medium. This degradation requires cells of the a/a mating type and nutrient sensing through the Snf1 protein kinase, and is mediated *in trans* by APC/C^Cdh1^. Cdh1-Cdh1 recruitment did not occur through recognition of a typical D-box or KEN box motif, but was inhibited by CDK-mediated phosphorylation within its N-terminal domain. Thus, Cdh1 degradation provides a novel example of APC/C self-regulation that redirects its enzymatic activity during nutrient starvation and preserves the accumulation of Cdh1 substrates needed for the completion of the meiotic program.

## RESULTS

### Transition to sporulation leads to turnover of Cdh1

During vegetative growth of *S. cerevisiae*, Cdh1 undergoes cycles of phosphorylation and de-phosphorylation and binding to regulatory proteins, but its steady-state protein levels remains relatively constant. We tested a range of growth conditions to determine if any affected the stability or steady-state level of Cdh1. The most dramatic effect was observed upon transfer of cells from rich medium (YPD) to sporulation medium (SPM), which induces cells to enter meiosis and to develop into spores as a protective mechanism during poor environmental conditions. The critical components of SPM are a very low concentration of sugar (0.02% raffinose vs the usual 2.0% glucose, 0.3% potassium acetate, and very low concentrations of amino acids). As shown in Figure 1A (*top panel*), Cdh1 levels remained relatively constant when cells were maintained in rich medium but dropped precipitously between six and twelve hours following the shift to sporulation medium.

**Figure 1.**
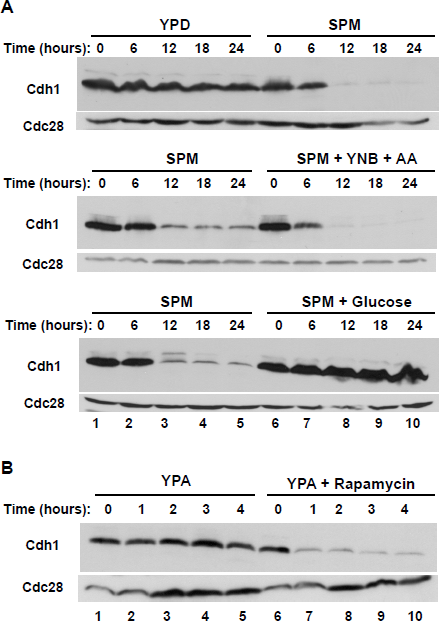
Cdh1 protein levels decline during sporulation. **(A)** *(top)* Cdh1 protein in extracts from an asynchronous population of diploid yeast cells (strain DOY2361) grown in rich medium (YPD, lanes 1-5) or cells transferred from YPD to sporulation medium (SPM, lanes 6-10). Samples were withdrawn at the indicated times and monitored for the presence of Myc-Cdh1 by immunoblotting with 9E10 antibodies. As a loading control, the membrane was re-probed with anti-PSTAIR antibodies to detect Cdc28. *(middle)* As above with cells transferred to sporulation medium (SPM, lanes 1-5) or to sporulation medium supplemented with 0.67% yeast nitrogen base, 0.2% amino acid mixture (SPM + YNB + AA, lanes 6-10). *(bottom)* As above with cells transferred to sporulation medium (SPM, lanes 1-5) or to sporulation medium supplemented with 1% glucose (lanes 6-10). **(B)** Asynchronous cultures of diploid yeast cells (strain DOY2361) pre-grown in acetate-containing medium (YPA) were mock-treated (lanes 1-5) or incubated with 1 µg/ml rapamycin (lanes 6-10). Samples were withdrawn at the indicated times and monitored for the presence of Cdh1 by immunoblotting with anti-Myc antibodies.

We explored which nutritional changes were needed for the reduction in Cdh1 level. The addition of amino acids and nitrogen sources to sporulation medium had little effect on Cdh1 levels (Figure 1A, *middle panel*), suggesting that nitrogen starvation per se was not a required signal. In contrast, addition of 1% glucose (or galactose (not shown)) fully prevented Cdh1 down-regulation (Figure 1A, *bottom panel*). The addition of glucose also prevented the transition to sporulation, which was evident by the absence of tetrads even after prolonged times. Thus, a glucose sensing pathway is critical not only for preventing the initiation of the sporulation program but also for the maintenance of Cdh1 protein expression and stability. We will explore the relationship between these events below (Figure 4).

**Figure 4.**
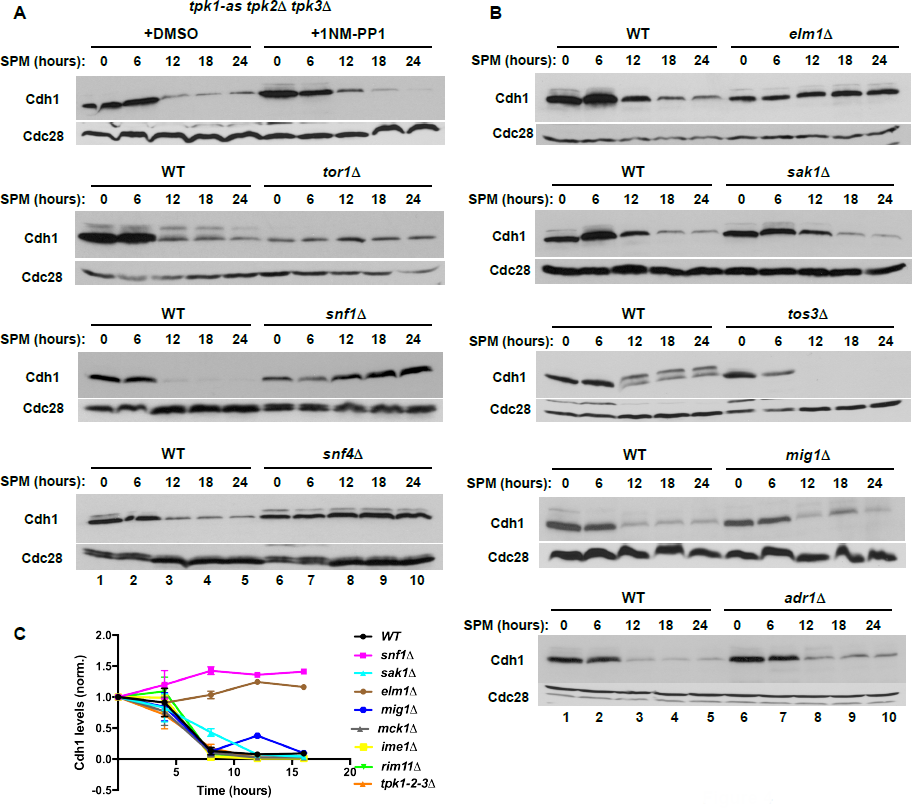
The glucose-sensing Snf1 protein kinase pathway is required for Cdh1 turnover. **(A)** *(top)* The cAMP-dependent protein kinase A (PKA) is not involved in Cdh1 turnover. Yeast PKA activity can be eliminated by deleting two of the three PKA isoforms (Tpk2, Tpk3) and expressing an analog-sensitive version of the third isoform (Tpk1, *tpk1-as*). Homozygous cells (strain DOY2596) carrying *tpk1-as tpk2Δ tpk3Δ* mutations were grown in YPD, transferred to sporulation medium in the absence (lanes 1-5) or presence (lanes 6-10) of 1 µg/ml of the 1NMPP1 nucleotide analog that rapidly inactivates Tpk1-as. *(panels two to four)* Wild-type (strain DOY2361) and isogenic mutant strains carrying homozygous deletions of genes implicated in glucose and nutrient signaling pathways - *TOR1* (strain DOY2512), *SNF1* (strain DOY2603), and *SNF4* (strain DOY2655) - were grown in YPD and transferred to sporulation medium as in Figure 1A. **(B)** *(panels one to three)* Wild type (strain DOY2361) and three isogenic mutant strains carrying deletions of Snf1-activating kinases, *ELM1* (strain DOY2676), *SAK1* (strain DOY2675), and *TOS3* (strain DOY3016) were grown in parallel and transferred to sporulation-inducing medium as in (A). Samples were withdrawn at the indicated times and probed to measure Cdh1 protein levels with 9E10 antibodies. *(panels four* and *five)* Mig1 and Adr1 are transcription factors that function downstream of Snf1. Wild-type and isogenic diploid strains carrying *mig1Δ* (strain DOY2853) and *adr1Δ* (strain DOY2651) deletions were tested in sporulation - inducing medium as in (A). Following transfer to SPM, samples were withdrawn at the indicated times and processed for immunoblotting with 9E10 antibodies to detect Myc-Cdh1. Anti-Cdc28 probing was used as a loading control. **(C)** The relative abundances of Cdh1 in wild-type and the indicated mutant strains were quantified by ImageJ software and normalized relative to the level of Cdc28 from the same sample. The initial amount of Cdh1 at time 0 was set as 100% and the fractions of the protein remaining in the samples at subsequent time points were plotted relative to this starting fraction. Some of the data are derived from experiments depicted in Figure 4S.

In addition to a nutrient-poor environment, sporulation can be triggered by pharmacological inhibition of the TOR signaling pathway, which mimics starvation for nitrogen and phosphate. Cells grown in YPA (rich in nitrogen sources and acetate, but lacking glucose) maintained their level of Cdh1 (Figure 1B). In contrast, addition of rapamycin caused a rapid (under one hour) degradation of Cdh1. Thus, cells will degrade Cdh1 if they think they’re starving for glucose and nitrogen, even when nitrogen is abundant. We suspect that the more rapid degradation of Cdh1 when cells are transferred from YPA to YPA + rapamycin than when they are transferred from YPD to SPM relates to internal nutrient supplies. Cells switching from YPD to SPM must deplete their internal stores of glucose and of excess amino acids. In contrast, cells switching from YPA to YPA + rapamycin have no glucose to begin with and are ‘tricked’ by the presence of rapamycin into thinking they have low nitrogen stores.

In addition to rapamycin, caffeine and methionine sulfoximine (MSX), two chemically unrelated compounds that also inhibit the TOR signaling pathway, also decreased Cdh1 levels, whereas other types of stress conditions, such as high osmolarity, heat shock, oxidative stress, and DNA damage had no detectable effect (data not shown).

### Diploid requirement for Cdh1 degradation

Because the experiments in Figure 1 used diploid *MATa/MAT*a cells, we wondered whether the degradation of Cdh1 in sporulation medium was a general response to starvation, or if it requires diploidy. Natural population of *S. cerevisiae* exist in three developmentally distinct mating types, which are distinguished by the expression of transcription factors from their mating-type loci. **a** and α haploids contain *MATa* or *MATα* at their mating-type loci, respectively whereas **a/α** diploids are heterozygous for this locus (*MATa/MATα*). Diploid cells can undergo meiosis and sporulation to produce haploid spores, which can germinate to produce haploid cells capable of mating with cells of the opposite mating type to re-form diploids.

Although Cdh1 was unstable in a/α diploids grown in sporulation medium (Figure 2), Cdh1 was fully stable in *MATa* haploids, which arrested as individual unbudded cells rather than proceeding though sporulation. Since **a/α** diploids differ from **a** haploids in both ploidy and mating type, we tested Cdh1 stability in **a/a** diploids to distinguish which factor was important. **a/a** diploids have the ploidy of **a/α** cells but have the mating type of **a** cells. Interestingly, Cdh1 was fully stable in **a/a** diploid cells shifted to sporulation medium (Figure 2, *middle panel*), indicating that it is mating type, and not ploidy, that is important for the decline in Cdh1 levels. Finally, we found that addition of the ***α2*** transcription factor to a *MATa* haploid, thereby converting these cells to the *MATa/MATα* transcriptional profile, led to Cdh1 turnover (Figure 2, *lower panel*). Therefore, Cdh1 down-regulation requires the transcription profile of *MATa/MATα* cells but does not require diploidy *per* se.

**Figure 2.**
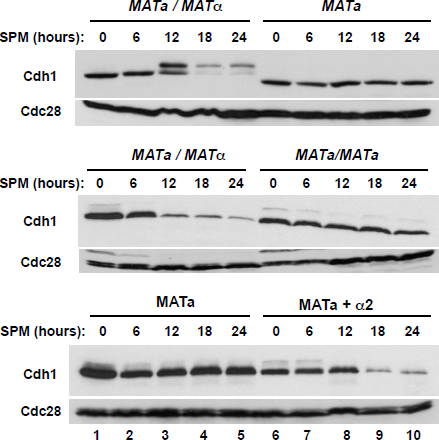
Cdh1 degradation depends on heterozygosity of the mating type locus but not on cell ploidy per se. *(top)* Asynchronous diploid **a / α** cells (strain DOY2361) (lanes 1-5) and haploid **a** cells (strain DOY2282) (lanes 6-10) were transferred from YPD to sporulation medium as in Figure 1A. Samples were withdrawn at the indicated times and processed for immunoblotting to detect Myc-tagged Cdh1. Anti-Cdc28 probing of the same filter was used as a loading control. *(middle)* The same time courses as above were performed using two isogenic diploid strains that differ only in the configuration at the mating-type locus: **a/**α (strain DOY2780) (lanes 1-5) and **a/α** (strain DOY2783) (lanes 6-10). Only the **a/**α cells were able to sporulate and produce tetrads. *(bottom)* The same time courses were performed using a haploid **a** strain carrying either an empty vector (lanes 1-5) or a vector with expressing the α**2** transcription factor (strain DOY2786) normally expressed in **a/**α diploid cells, thus making these cells equivalent to **a/**α cells at the mating-type locus but otherwise haploid (lanes 6-10). The **a** *+* α**2** cells could not undergo sporulation.

### The initiation of meiosis is not required for Cdh1 down-regulation

Although we have shown that conditions that lead to meiosis and sporulation (diploidy and starvation) also lead to reduced levels of Cdh1, we have not shown that entry into meiosis is necessary for the decline in Cdh1 level. To do so, we examined Cdh1 levels in three *IME* (*I*nitiator of *ME*iosis) mutants. *IME1* encodes the major transcriptional activator of early meiotic genes. Its promoter harbors numerous cis-acting sensors for cell ploidy, glucose, and nitrogen that together determine whether external conditions are appropriate to begin the transition into meiosis and sporulation. *IME2* encodes a Cdc28-like protein kinase that phosphorylates multiple targets, including Cdh1, and is a direct transcriptional target of Ime1. *IME4* encodes an mRNA adenosine methyltransferase that acts independently from *IME1* and *IME2* to stimulate expression of early meiotic genes. In the absence of any one of these genes, cells fail to enter the meiotic program and continue in the vegetative state. Remarkably, deletion of any of these meiotic inducers had no effect on the decline of Cdh1 (Figure 3). Thus, the down-regulation of Cdh1 is signaled prior to or in parallel to the induction of meiotic entry by the *IME* genes.

**Figure 3.**
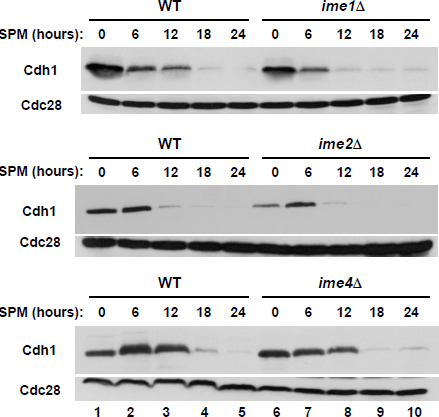
Expression of **I**nducer of **ME**iosis genes is not required for Cdh1 degradation. Homozygous diploid strains that were wild-type (strain DOY2361) or deleted for *IME1* (strain DOY2554), *IME2* (strain DOY2555), or *IME4* (strain DOY2977) were grown in YPD medium, transferred to sporulation medium, and tested at the indicated times for Cdh1 expression. Mutation of each *IME* gene prevents transcription of early meiotic genes and sporulation. Samples were processed for immunoblotting with 9E10 antibodies to detect Myc-Cdh1 protein. Cdc28 was used as a loading control.

### Cdh1 down-regulation is mediated by the *SNF1* protein kinase complex

One of the major pathways for sensing glucose availability in both yeast and higher eukaryotes is through protein kinase A (PKA). Glucose starvation inhibits PKA activity and PKA inhibits the sporulation process. Thus, glucose starvation induces the sporulation pathway. If PKA controls Cdh1 stability, then inhibition of PKA should prevent the decline in Cdh1 level, even in the absence of glucose. As in higher eukaryotes, yeast PKA is composed of two regulatory and two catalytic subunits, the later encoded by three closely related genes - *TPK1, TPK2* and *TPK3*. Although these catalytic subunits are redundant, at least one is necessary for survival. To eliminate PKA activity conditionally, we used a yeast strain deleted for genes for two of the catalytic subunits and encoding an ‘analog-sensitive’ version of the third so that it could be chemically inhibited. Cdh1 levels declined whether or not this *tpk1-as tpk2Δtpk3Δ* strain was treated with the ATP analog, 1NM-PP1, to inhibit Tpk1-as (Figure 4A, *top panel*). Thus, PKA was unlikely to control Cdh1 stability.

The TOR pathway is another major nutrient-sensing pathway that converges to impact the transition to sporulation. TORC1 is a conserved protein kinase complex implicated in cellular sensing of essential nutrients, including nitrogen, phosphate, and amino acids. During nitrogen starvation, TORC1 kinase activity is suppressed, which leads to cellular commitment to initiate the sporulation process. Furthermore, pharmacological inhibition of TORC1 kinase activity activates the transcription of early meiotic genes and triggers the transition to sporulation even in the presence of abundant nutrients. Since the TORC1 inhibitor rapamycin accelerated Cdh1 turnover, we asked whether Tor1, the catalytic subunit of the complex, was required for Cdh1 stability. Although the basal expression of Cdh1 in *tor1Δ* cells was significantly lower than in wild-type cells, Cdh1 levels did not change significantly upon transfer to sporulation medium (Figure 4A, *second panel*). From both this experiment and the rapamycin inhibition experiment in Figure 1B, it appears that TORC1 exerts a protective effect on Cdh1 stability, but that it is not the primary target for the destabilization of Cdh1 upon glucose withdrawal.

In a similar manner, we found that two other proteins involved in nutrient signaling and the initiation of meiosis and sporulation (Rim11 and Mck1) also had no impact on Cdh1 down-regulation Figure 4S).

A major pathway for sensing the presence of glucose involves the SNF1 protein kinase complex. The SNF1complex (ortholog of the mammalian AMPK nutrient sensor) is required for yeast cells to survive in the absence of fermentable glucose and other environmental stresses. SNF1 protein kinase activity increases in the presence of alternative carbon sources and during sporulation but is rapidly inhibited once glucose becomes available. In the absence of Snf1, SNF1-regulated pathways are unaffected by the absence of glucose. Notably, deletion of *SNF1* prevented the down-regulation of Cdh1 in sporulation medium (Figure 4A, *third panel*, lanes 6-10). In addition to Snf1, the SNF1 complex contains one of three alternative β subunits, which differ in their subcellular localizations and substrate specificity, and a unique γ-subunit, Snf4. None of the single β subunit deletions (*GAL83, SIP1,* and *SIP2*) affected Cdh1 levels (Figure 4S), indicating functional redundancy among these proteins. In contrast, deletion of the sole γ-subunit, *SNF4*, fully stabilized Cdh1 (Figure 4A, *forth panel*).

In addition to binding Snf4, activation of Snf1 kinase requires phosphorylation at Thr270 by one of three upstream protein kinases: Sak1, Elm1, and Tos3. The activities of these partially redundant protein kinases are ultimately linked to intracellular AMP concentration, which in turn depends on the availability of glucose. We tested whether deletion of any of these upstream protein kinases affected Cdh1 levels. Surprisingly, deletion of *SAK1* had negligible effects on Cdh1 stability (Figure 4B, *second panel*), even though its abundance and overall activity are significantly higher than those of the other two activating kinases (Ben Turk, Yale University, personal communication). Deletion of *TOS3* also had no effect on Cdh1 levels. However, deletion of *ELM1* fully stabilized Cdh1, resembling the effects observed in *snf1Δ* and *snf4Δ* mutant strains (Figure 4B, *top panel*). Thus, of the three protein kinases directly upstream of Snf1, Elm1 acts uniquely to activate Snf1 in a pathway from glucose limitation to Cdh1 down-regulation. However, it should be noted, that in addition to Snf1, Elm1 phosphorylates numerous other protein targets, so we cannot exclude the possibility that the stabilizing effect of *elm1Δ* might occur through another pathway.

SNF1 exerts its major effects through large-scale reprogramming of gene transcription in response to glucose depletion. For example, Snf1 directly phosphorylates the Mig1 transcriptional repressor, which leads to its release from gene promoters required for utilization of alternative carbon sources. Snf1 also phosphorylates the transcriptional activator Adr1, leading to the transcription of many metabolic genes, and the N-terminal domain of histone H3, which leads to transcriptional changes within hundreds of genes. To examine the possibility that increased Cdh1 turnover is caused by changes in gene transcription downstream of Snf1, we examined Cdh1 protein levels in *mig1Δ* and *adr1Δ* mutant strains. We found that neither deletion affects Cdh1 levels–in sharp contrast to the effect in *snf1Δ* and *snf4Δ* cells-making it unlikely that expression of one of the many Mig1-repressed or Adr1-activated genes is responsible for Cdh1 instability (Figure 4B, *bottom panels*).

### APC/C-dependent degradation of Cdh1

Before determining which ubiquitin pathway components might be responsible for the decline in Cdh1 level, we first verified that the decrease in Cdh1 is indeed ubiquitin-dependent. Addition of the proteasome inhibitor MG-132 blocked the decline in Cdh1 upon transfer to sporulation medium (Figure 5A). The gradual increase in the amount of Cdh1 is presumably due to new synthesis in the absence of degradation, similar to that observed upon deletion of *SNF1* (Figure 4A).

**Figure 5.**
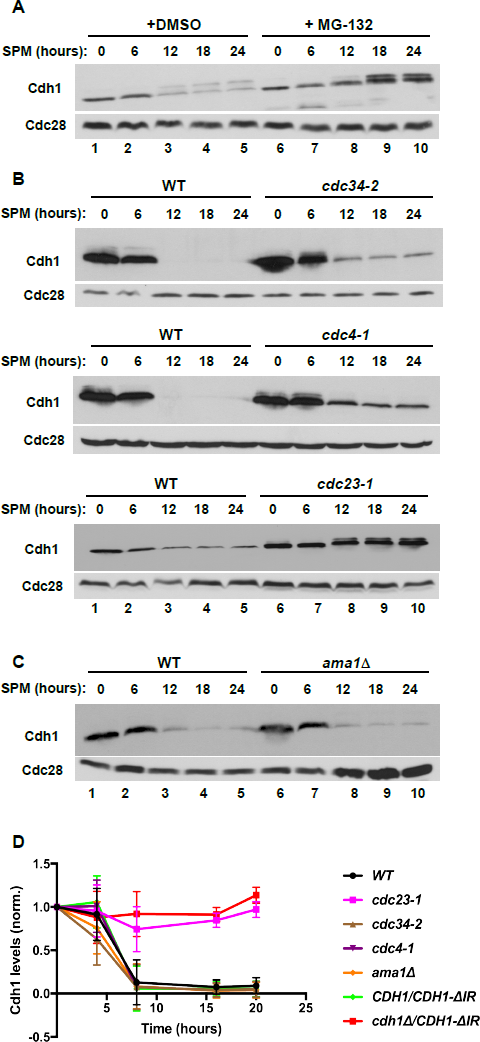
Cdh1 is degraded in an APC/C-dependent manner during sporulation. **(A)** *pdr5Δ* homozygous diploid cells (which are permeable to the proteasome inhibitor MG132; strain DOY2989) were transferred to sporulation medium in the absence (lanes 1-5) or presence of 20 µM MG132 to inhibit the proteasome. Samples were withdrawn at the indicated times and processed for immunoblotting with 9E10 antibodies to detect Myc-Cdh1 as in Figure 1A. Anti-Cdc28 was used as a loading control. **(B)** Wild-type and isogenic homozygous diploid strains carrying a conditional allele of the ubiquitin-conjugating enzyme of the SCF complex (*cdc34-2*; strain DOY2542), the F-box protein adaptor of the SCF (*cdc4-1*; strain DOY2541), and an APC/C core subunit (*cdc23-1*; strain DOY2540) were grown in YPD medium at permissive conditions and transferred to sporulation medium at 35°C to inactivate the temperature-sensitive proteins. Samples were withdrawn at the indicated times and processed to visualize Myc-Cdh1 by immunoblotting with 9E10 antibodies. Of note, transfer to 35°C did not interfere with the ability of wild-type cells to form tetrads but arrested cell cycle progression of the *cdc34-2, cdc4-1* and *cdc23-1* strains. **(C)** Wild-type cells and an *ama1Δ* mutant strain (strain DOY2602), which carries a homozygous deletion of the meiosis-specific APC/C activator, were transferred to sporulation-inducing medium and incubated at 30°C for the indicated times. Samples were processed to measure Cdh1 protein levels as in Figure 1A. **(D)** The relative stability of Cdh1 in wild-type and the indicated mutant strains was quantified by ImageJ software. The initial level of Cdh1 in each individual strain at time 0 was set to 100%. The data for *CDH1-ΔIR* strains comes from experiments shown in Fig. 6. Data represent the means plus standard deviations from at least three independent experiments.

Since the Snf1 protein kinase is required for Cdh1 degradation and since substrates are often targeted to the SCF ubiquitin ligase via phosphorylation, we turned our attention to the possible involvement of an Skp1-Cullin-F box complex. We shifted wild-type or temperature-sensitive mutant strains into sporulation medium at the non-permissive temperature (35°C) for the mutants and followed Cdh1 levels. Cdc34 is a ubiquitin-conjugating enzyme (E2) that functions with all SCF complexes in yeast. Cdc4 is a major F-box protein for the SCF. We noticed that wild-type cells were able to sporulate and form tetrads at the elevated temperature (35°C), whereas *cdc34-2* mutant arrested as single cells, likely due to abnormally high levels of multiple SCF substrates. Even though *cdc34-2* cells did not proceed through meiosis, Cdh1 was still down-regulated to the same extent as in wild-type cells (Figure 5B, *top panel*). Similarly, inactivation of Cdc4 also failed to stabilize Cdh1 (Figure 5B, *middle* panel).

We next asked whether the APC/C itself might promote Cdh1 degradation. We envisioned that Cdh1 might transition from being an APC/C activator to being an APC/C substrate. Two examples of such a transition are well documented in vegetative cells, first, in which Cdc20 within an APC/C^Cdc20^ complex ‘auto-ubiquitinates’ (‘*in cis*’) during mitosis, and second, in which APC/C^Cdh1^ promotes the ubiquitination of Cdc20 (*‘in tran’s*) in G1. We tested whether the APC/C was involved in Cdh1 degradation by transferring wild-type cells and *cdc23-1* cells (containing a temperature-sensitive mutation in a core APC/C subunit) into sporulation medium at the non-permissive temperature (35°C) and following Cdh1 levels. The elevated temperature did not affect Cdh1 degradation in wild-type cells but it completely blocked degradation in the *cdc23-1* cells (Figure 5B, *bottom panel*), indicating that the APC/C is involved in Cdh1 degradation.

### Autoubiquitination of Cdh1 is mediated by APC/C^Cdh1^ *in trans*

Having determined that the APC/C core complex was essential for Cdh1 turnover, we next asked which activator protein was involved in this process. Ama1 is a meiosis-specific homolog of Cdc20 and Cdh1 that promotes the ubiquitination of multiple regulatory proteins, including Cdc20. Since *AMA1* is not essential for vegetative growth, we examined Cdh1 levels in *ama1Δ* mutant cells after their transition to sporulation medium. As expected, *ama1Δ* cells were unable to enter meiosis. However, Cdh1 turnover occurred as well in *ama1Δ* cells as in wild-type cells (Figure 5C), indicating that Ama1 was not required for Cdh1 turnover.

Excluding Ama1 raised the possibility that Cdh1 might promote its own degradation. One version of such auto-ubiquitination (termed ‘*trans*’) involves two molecules of Cdh1, one occupying the conventional activator position in an APC/C^Cdh1^ complex, and the second serving as the substrate that is bound to the APC/C^Cdh1^ complex. To test this possibility, we expressed a form of Cdh1 termed Cdh1-ΔIR, which lacks its two C-terminal amino acids and is thus unable to interact with the APC/C core subunit Cdc23. It is thus unable to serve as an APC/C activator, but could potentially serve as a substrate. (Preliminary testing in haploid cells showed that Cdh1-ΔIR was unable to support the degradation of the Clb2 APC/C substrate (data not shown).) When cells expressing Cdh1-ΔIR but no other form of Cdh1 were transferred to sporulation medium, Cdh1-ΔIR was stable (Figure 6A, *top panel*, lanes 6-10). However, when the cells also expressed wild-type Cdh1, Cdh1-ΔIR was degraded with normal kinetics (Figure 6A, *top panel*, lanes 1-5), indicating that it was degraded *in trans*.

**Figure 6.**
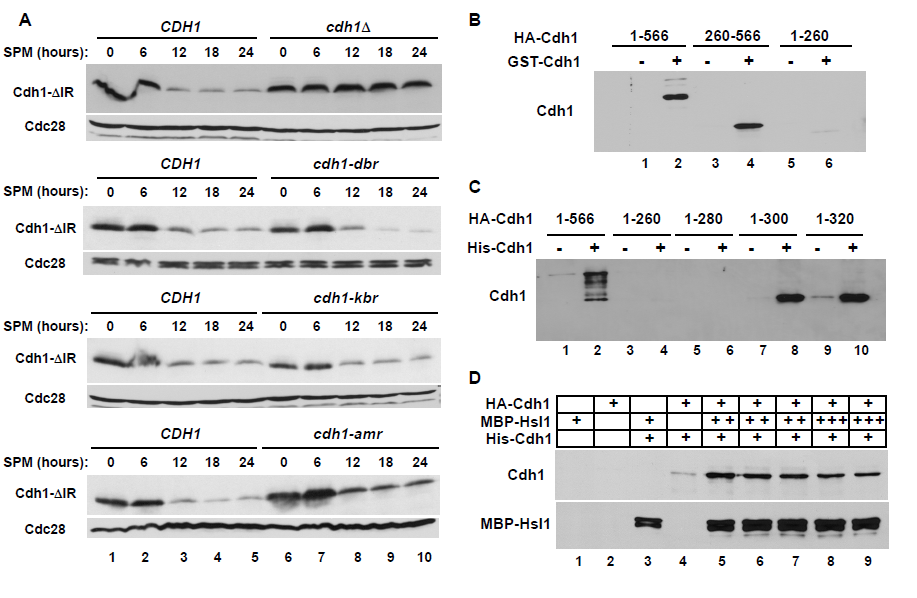
Atypical *trans* degradation of Cdh1 by APC/C^Cdh1^. **(A)** All strains contain *Myc-CDH1-IRΔ* (‘Cdh1-ΔIR’), which lacks the two C-terminal amino acid residues of Cdh1 and is thus unable to bind to the APC/C core. Therefore, it is non-functional for promoting the degradation of APC/C substrates. As indicated, each strain also contained a wild-type copy of *CDH1* (strain DOY2719), an empty vector (*cdh1Δ,* strain DOY2725), or a form of Cdh1 that has been mutated so that it can no longer recognize Destruction Boxes (*cdh1-dbr*; strain DOY2971), KEN boxes (*cdh1-mkr*; strain DOY2972), or the ABBA motif (*cdh1-amr*; strain DOY2973). The resulting strains were grown in YPD and transferred to sporulation medium. Samples were withdrawn at the indicated times and immunoblotted with 9E10 antibodies to detect Myc-Cdh1-ΔIR. Anti-Cdc28 probing was used as a loading control. **(B)** Wild-type GST-Cdh1 was expressed and purified from yeast cells. Full-length HA-Cdh1 (lanes 1-2), a fragment corresponding to the Cdh1 WD-40 domain (amino acids 260-566, lanes 3-4), and the N-terminal portion of Cdh1 (amino acids 1-260, lanes 5-6), were expressed by translation *in vitro*, incubated with control beads (lanes 1, 3, and 5) or GST-Cdh1 beads (lanes 2, 4, and 6) and bound proteins were analyzed by immunoblotting with 12CA5 antibodies to detect HA-Cdh1. **(C)** HA-Cdh1 (amino acids 1-566 lanes 1-2), and its indicated fragments (lanes 3-10) were translated *in vitro*, incubated with control and His_6_-Cdh1 beads, and bound proteins were visualized by immunoblotting to detect HA-Cdh1 fragments as in (A). **(D)** Full-length *in vitro* translated HA-Cdh1 was incubated with control (lane 2) and His_6_-Cdh1 beads in the absence (lane 4) and presence of increasing amounts of recombinant MBP-Hsl1 protein (lanes 5-9). Proteins associated with His_6_-Cdh1 were analyzed by immunoblotting with anti-HA and anti-MBP antibodies.

Although most APC/C substrates and other Cdc20- or Cdh1-binding proteins contain one of the common degradation motifs (a Destruction Box, a KEN box, or, less commonly, an ABBA motif), Cdh1 itself contains none of these motifs. Some APC/C substrates contain a degenerate form of one of these motifs that binds to the same binding surface on Cdc20 or Cdh1 as conventional substrates. Structural studies have revealed the ‘receptors’ within the WD-40 domains of the APC/C activators that are specific for the three different types of APC/C degrons. We determined whether Cdh1 might contain a ‘cryptic’ degradation motif by testing whether Cdh1-ΔIR was stabilized in the presence of otherwise wild-type Cdh1 containing one of the mutant receptors. Remarkably, Cdh1-ΔIR was degraded with wild-type kinetics in the presence of Cdh1 with a Destruction Box receptor (*cdh1-dbr*), a KEN box receptor (*cdh1-dkr*), or an ABBA receptor (*cdh1-amr*) (Figure 6A, *panels two to four*) even though these mutants were deficient in targeting known APC/C substrates in G1 cells (Qin *et al*., 2016). Thus, Cdh1-Cdh1 recognition during the sporulation program involves a non-canonical degradation motif on the substrate Cdh1 and a novel binding surface on the activator Cdh1.

### Direct Cdh1-Cdh1 binding

We examined whether Cdh1 could bind to itself *in vitro*. We immobilized full-length wild-type Cdh1 on beads that were then incubated with various forms of N-terminally tagged Cdh1 that were translated *in vitro*. In a series of pull down experiments, we found that soluble full-length (1-566) Cdh1 specifically interacted with immobilized Cdh1, indicating that the two proteins could form at least a dimer (Figures 6B and 6C). We also observed similar Cdh1-Cdh1 interactions with proteins isolated from yeast extracts by immunoprecipitation (data not shown). By testing truncation mutations that were translated *in vitro*, we found that the C-terminal WD-40 domain was not necessary for binding to a full-length activator (Figure 6B and 6C). Interestingly, Cdh1 fragment 1-300 aa was a strong binder, whereas a slightly shorter fragment (1-280 aa), was unable to form a complex, suggesting that amino acids 280-300 may be critical for the Cdh1-Cdh1 interaction (Figure 6C).

We probed the nature of the Cdh1-Cdh1 interaction by performing binding reactions in the presence of a competitive inhibitor, MBP-Hsl1. Hsl1 is one of the best-characterized APC/C substrates and contains typical D-box and KEN motifs with a strong affinity to Cdh1. As expected, Hsl1 bound efficiently Cdh1 on beads. Interestingly, the presence of even high and beyond saturating levels of MBP-Hsl1 had little effect on the amount of Cdh1 bound (Figure 6D), consistent with Cdh1 binding to a non-canonical binding surface on the bead-bound Cdh1.

### CDK-mediated phosphorylation may regulate Cdh1 stability

Because the N-terminal half of Cdh1 is highly unstructured, charged, and subject to phosphorylation by multiple protein kinases, we suspected this region might contribute to or regulate Cdh1 degradation. We constructed a set of N-terminal deletions using a tagged copy of Cdh1 and tested these substrates *in vivo* using a panel of yeast strains carrying an untagged wild-type copy of Cdh1 (Figure 7). The resulting heterozygous strains were exposed to sporulation conditions and analyzed for their ability to degrade Cdh1 ‘substrate’ proteins. Deletion of the first fifty amino acids, which removed the NLS and C-box (important for APC/C binding), had no effect on Cdh1 stability (Figure 7, *upper panel*). Removing 150 amino acids significantly stabilized the Cdh1 (Figure 7, *third panel*), as did an internal deletion of amino acids 50-157 (Figure 7, *bottom panel*). The results of testing other deletions were consistent with these findings, but were unable to pinpoint the degron more specifically or to identify key conserved residues responsible for Cdh1 instability. We do not know if this region contains a degron for Cdh1 degradation, or if its proper folding is necessary for the presentation of a degron located elsewhere.

**Figure 7.**
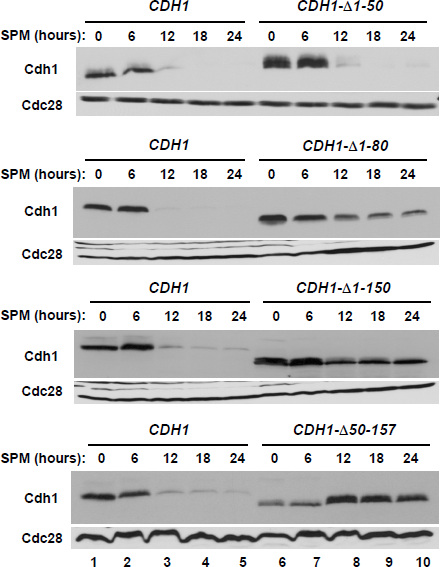
The N-terminal domain of Cdh1 contains a degradation motif. Endogenous wild-type *Myc-CDH1* and the indicated truncations were co-expressed with an untagged copy of *CDH1* in *MATa/MATα* diploid strains. The resulting heterozygous strains (*Myc-CDH1*/*CDH1* (strain DOY2361), *Myc-CDH1-Δ1-50*/*CDH1* (strain DOY2500)*, Myc-CDH1-Δ1-80*/*CDH1* (strain DOY2476)*, Myc-CDH1-Δ1-150*/*CDH1* (strain DOY2477)*, and Myc-CDH1-Δ50-157*/*CDH1)* (strain DOY2834) were tested for Myc-Cdh1 stability after transition to sporulation medium. Yeast extracts prepared from the samples withdrawn at the indicated time points were immunoblotted with anti-Myc antibodies to detect Cdh1. The membranes were then re-probed with anti-Cdc28 antibodies to serve as a loading control.

A number of protein kinases, including Cdc28 (Cdk1), Cdc5 and Ime2 have been found to phosphorylate many residues within the Cdh1 N-terminal region. For example, Cdc28 phosphorylates Cdh1 on nine distinct sites that undergo cycles of phosphorylation/dephosphorylation during the mitotic cell cycle. These phosphorylations regulate both Cdh1 localization and its ability to bind APC/C. These Cdc28 sites can be subdivided into two groups, according to their location and functionality (Figure 8A) (Hockner *et al*., 2016): sites 1-3 regulate Cdh1 localization, whereas sites 4-9 mediate its interaction with APC/C. Dephosphorylation of Cdh1 sites 1-3 leads to preferential accumulation of Cdh1in the nucleus. We found that mutations of sites 1-3 to alanine or to aspartic acid had no significant effect on Cdh1 stability in sporulation medium (Figure 8B, *upper panel*), which is consistent with the lack of effect of deletion of the NLS sequence (Figure 7, *top panel*). Thus, nuclear accumulation or cytoplasmic retention appear to have no significant impact on Cdh1 turnover in sporulation conditions. In sharp contrast, mutations of sites 4-9 or even 4-7 to aspartic acid resulted in a profound increase in Cdh1 stability (Figure 8C, *middle* and *bottom panels*). This reversal of protein turnover was remarkably similar to that observed during glucose inhibition and in *snf1*Δ mutant yeast strains (compare Figures 4 and 8). Thus, it appears that putative phosphorylation at sites 4-7 protects Cdh1 from APC/C^Cdh1^-mediated ubiquitination following transfer to sporulation medium, when Cdh1 would normally be unstable.

**Figure 8.**
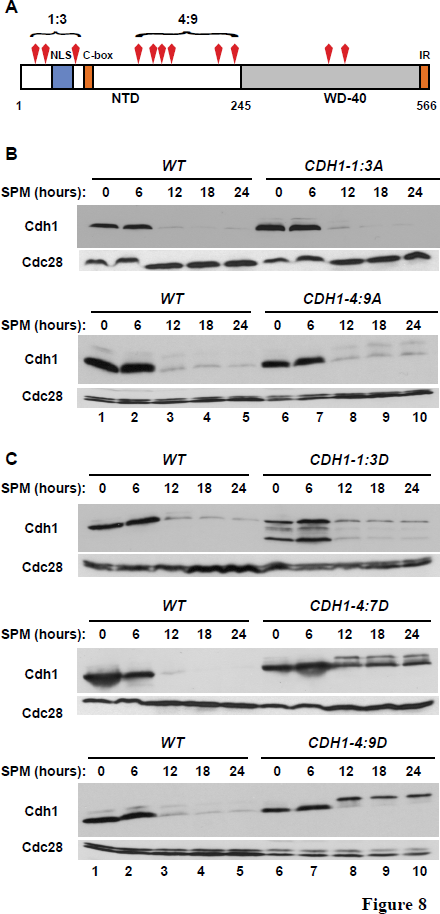
Putative Cdc28-mediated phosphorylation regulates Cdh1 stability in sporulation medium. **(A)** A diagram of Cdh1 indicating the locations of CDK phosphorylation sites and of subsets of these sites within the N-terminal domain used in the various mutants. The first cluster of CDK sites, Cdh1-1-3, (Thr12, Ser16, and Ser42) is located near the NLS motif and controls protein localization. The second cluster of CDK sites, Cdh1-4-9, (Ser157, Ser169, Thr173, Thr176, Ser227, Ser239) regulates Cdh1 interaction with the core APC. The locations of the C-box and the IR motif are indicated. **(B)** Two clusters of CDK sites within the Cdh1 N-terminal domain (A) were mutated to alanine. Both *CDH1-1-3A* and *CDH1-4-9A* were co-expressed with an untagged copy of wild-type *CDH1* in *MATa/MATα* diploid strains. The various forms of Cdh1 were tested for their stability upon shift to sporulation medium as in Figure 1A. **(C)** Clusters of the same nine CDK sites were mutated to aspartic acid to mimic phosphorylation. *CDH1-1:3D*, *CDH1-4:9D* were as indicated in (A); *CDH1-4:7D* carried S157D, S169D, T173D, T176D. These mutant alleles of *CDH1* were co-expressed with untagged *CDH1* in *MATa/MATα* diploid strains. The various forms of Cdh1 were tested for their stability upon shift to sporulation medium as above.

## DISCUSSION

The activities of key cell cycle regulators, such as APC/C^Cdh1^, in yeast and in higher eukaryotes are tightly regulated through the integration of signals from internal and external sources. We have discovered a complex network of signals leading to the abrupt degradation of Cdh1 during the transition from vegetative growth to the meiotic/sporulation program. Destabilization of Cdh1 required heterozygosity at the mating-type locus (**a/**α cells) and starvation for glucose. The apparent degradation of Cdh1 was mediated by the 26S proteasome and involved ‘auto-ubiquitination’ of Cdh1 by APC/C^Cdh1^ operating *in trans*.

There are striking similarities and differences in the regulation of Cdh1 levels by phosphorylation in human and yeast cells. In both species, the Cdh1 N-terminal domain is mostly unstructured and contains multiple sites of phosphorylation regulating protein-protein interactions and protein stability. Several of these sites are evolutionary conserved, as are some of the protein kinases responsible for Cdh1 phosphorylation. The most prominent such protein kinase is CDK, which phosphorylates at least nine distinct sites within the N-terminal region of yeast Cdh1, thereby controlling protein localization and its binding to the core APC/C subunits. In human cells, cyclin A-CDK phosphorylation also provides a priming site for subsequent phosphorylation by Plk1, which is followed by β-TRCP binding and ubiquitination of Cdh1 by the SCF ^β-TRCP^ ubiquitin ligase (Lukas *et al*., 1999; Sorensen *et al*., 2000; Fukushima *et al*., 2013). In addition, cyclin F, another SCF adaptor protein, targets Cdh1 for SCF-mediated ubiquitination during early S-phase (Choudhury *et al*., 2016). Given the striking conservation of the yeast and human systems, we were surprised by the lack of any involvement of SCF complexes in the degradation of Cdh1 during sporulation. Rather, degradation appears to be totally dependent on the APC/C. Thus, even though Cdh1 was phosphorylated by conserved protein kinases, these phosphorylations serve to stabilize Cdh1 via an unknown mechanism rather than to target it for degradation mediated by an SCF complex.

The mechanism of Cdh1 degradation via the APC/C was surprising and quite different from what has been observed for other APC/C substrates. First, yeast Cdh1 does not contain any classical D-box, KEN box or ABBA motifs. Deletion analysis failed to identify any small region necessary for degradation. Second, we found that Cdh1 degradation occurred in *trans-*, i.e., it was mediated by a second functional Cdh1 molecule. This Cdh1-Cdh1 interaction failed to utilize any of the known substrate-recognition surfaces located within the C-terminal WD-40 domain of Cdh1-the D-box receptor, the KEN box receptor, or the ABBA receptor. Combined with the lack of competition between Hsl1 and Cdh1 for binding to Cdh1 on beads, these results suggest that the Cdh1-Cdh1 interaction involves a novel motif in the ‘substrate’ Cdh1 and a novel binding surface in the adapter’ Cdh1 molecule. Third, phospho-mimicking mutations within four sites-Ser157D, Set173D, Ser176D, Ser227D-stabilized Cdh1 during meiosis, suggesting that CDK-mediated phosphorylation may play a protective role to counteract APC/C-mediated ubiquitination. Phosphorylation-dependent stabilization has been reported for just a few APC/C substrates, including Pds1, Mps1, Fin1, and Nrm1 (Jaspersen *et al*., 2004; Woodbury and Morgan, 2007; Holt *et al*., 2008; Ostapenko and Solomon, 2011).

We can only speculate as to why Cdh1 needs to be degraded as cells transition to meiosis. In vegetative cells, for comparison, several overlapping mechanisms ensure tight cell cycle regulation of Cdh1 activity. Eliminating these controls is detrimental and leads to premature degradation of APC/C substrates. The key feature of meiosis-a reductional division followed immediately by an equational division without intervening Gap or S phases-requires that many proteins that would normally be degraded at the end of M phase persist. Such proteins include cyclins as well as regulators of chromosome cohesion. Specific APC/C substrates that might need to be preserved include Clb1 and Clb3, Cdc5 (polo-like kinase), Cdc20 and Sgo1(shugoshin). We suggest that in order to ensure rapid accumulation and maintenance of such proteins cells mark Cdh1 for self-degradation. Thus, cells re-program their APC/C activity to be more suitable for the meiotic program. This re-programing involves elimination of Cdh1 and the expression of the meiosis-specific APC/C adaptor Ama1.

It is interesting that the degradation of Cdh1, which seems necessary for proper meiosis and sporulation, doesn’t actually require that cells initiate the meiotic program. Since Cdh1 degradation occurs even in *ime1Δ* cells, it seems that the mating type and nutritional signals that tell cells to enter the meiotic program independently signal the degradation of Cdh1. Thus, in searching for proteins involved in the decision to degrade Cdh1, we can at once exclude all early, middle and late meiosis-specific gene products. Since cells enter into meiosis upon glucose and nitrogen limitation, it was not entirely surprising that inactivation of SNF1 activity was required for Cdh1 degradation. In fact, the SNF1 protein kinase complex acts independently of Ime1, and so is well positioned to be part of a signaling pathway leading to Cdh1 instability. In the presence of limited glucose and other environmental stresses, Snf1 is phosphorylated within its activation loop by one of three protein kinases-Sak1, Tos3, and Elm1 (Hong *et al*., 2003; Nath *et al*., 2003; Sutherland *et al*., 2003). It is unclear why Elm1 seems to play a predominant role in the signal to degrade Cdh1. We do not know how SNF1 signals Cdh1 degradation. One possibility is that it directly phosphorylates Cdh1 to causes its degradation. Alternatively, it could act through another factor. Mig1 and Adr1 are two direct targets of Snf1 that regulate the transcription of hundreds of glucose-sensitive genes (Treitel *et al*., 1998; Young *et al*., 2002). However, neither was necessary for the down-regulation of Cdh1. Thus, identifying the relationship between SNF1 and Cdh1 awaits further investigation. Similarly, we do not yet understand why Cdh1 degradation occurs in **a/**α cells but not in either **a** or α cells.

## MATERIALS AND METHODS

### Yeast strains and plasmids

Yeast strains were derivatives of W303a and W303a (*ade2-1trp1-1leu2-3,112 his3-11,15 ura3-1*); their relevant genotypes are listed in Supplementary Table 1. The *Myc_9_-CDH1* plasmid was provided by Wolfgang Seufert (University of Stuttgart, Stuttgart, Germany) (Zachariae *et al*., 1998). Conditional *cdc23-1, cdc28-13*, and *cdc34-2* strains were described previously (Burton and Solomon, 2000). The *MATa/MATa* diploid strain and *MATalpha2* expression vector were provided by Mark Hochstrasser (Yale University, New Haven, CT) (Hickey and Hochstrasser, 2015). The *tpk1-as tpk2Δtpk3Δ* strain was a generous gift of Angelika Amon (MIT, Cambridge, MA) (Weidberg *et al*., 2016). The *cdh1-dbr, cdh1-kbr, cdh1-amr* strains were provided by Mark Hall (Purdue University, West Lafayette, IN) (Qin *et al*., 2016). The *Myc_9_-CDH1 ime1Δ* (W303 *ime1::natMX4*), *ime2Δ* (W303 *ime2::natMX4*), *ime4Δ* (W303 *ime4::natMX4*), *snf1Δ* (W303 *snf1::natMX4*), *snf4Δ* (W303 *snf4::natMX4*), *sip1Δ* (W303 *sip1::natMX4*), *sip2Δ* (W303 *sip2::natMX4*), *gal83Δ* (W303 *gal83::natMX4*), *elm1Δ* (W303 *elm1::natMX4*), *mig1Δ* (W303 *mig1::natMX4*), *adr1Δ* (W303 *adr1::natMX4*), *tor1Δ* (W303a *tor1::natMX4*), and *sak1Δ* (W303a *sak1:: natMX4*) strains were accomplished by a PCR-based method (Goldstein and McCusker, 1999). Gene disruptions were verified by PCR using a primer downstream of the deleted gene and a primer internal to *natMX4*.

The *Myc_9_-CDH1* expression plasmid was used as template to introduce alanine or aspartate mutations within two clusters of CDK phospho-acceptor sites: T12, S16, S42 (*CDH1-1-3A* and *CDH1-1-3D*) and S157, S169, T173, T176, S227, S239 (*CDH1-4-9A* and *CDH1-4-9D*). The following in-frame deletions were introduced: *CDH1-Δ1-50, CDH1-Δ1-80, CDH1-Δ1-150, CDH1-Δ50-157*. The *GST-CDH1* expression plasmid was described previously (Burton and Solomon, 2000). All mutations were verified by sequencing of the entire coding region. Primer sequences and further details of the plasmids are available upon request.

### Cell growth and sporulation conditions

Cultures were grown in YPD, YP-Acetate (1% yeast extract, 2% peptone, 2% potassium acetate), and in complete minimal (CM) media as described (Guthrie and Fink, 1991). For sporulation time courses, cells were grown in YP-Acetate or YPD to mid-exponential phase (OD_600_ ˜0.4), washed by filtration (Corning Filter System), and released into sporulation medium (SPM) containing 0.3% potassium acetate, 0.02% raffinose, 20 µg/ml mixture of essential amino acids, adenine, and uracil for 24 hours at 30°C. Alternatively, conditional mutant *cdc23-1*, *cdc34-2*, and *cdc4-1* cells were grown at 23°C in YPD and released into SPM at 35°C for 24 hours. Starting from the release, equal volumes of cells (8 ml) were collected at 6-hour intervals by centrifugation at 4000 rpm for 2 min and frozen in liquid nitrogen.

For analyses of protein stability, haploid *MATa cdc28-13* cells were grown in YP-Raffinose to mid-exponential phase (OD_600_ ˜0.4), induced with 2% galactose for 50 min at 30°C, which resulted in protein synthesis at levels comparable to that of the endogenous protein, followed by addition of cycloheximide (500 µg/ml, MP Biomedicals) and 2% dextrose. Subsequently, equal volumes of cells (8 ml) were collected at 30-min intervals by centrifugation at 5000 rpm for 2 min and frozen in liquid nitrogen.

### Yeast extracts and Immunoblotting

Cells from 8 ml cultures (OD_600_ ˜0.4) were collected, washed with ice-cold H_2_0, suspended in 0.5 ml 100 mM NaOH and incubated for 5 min at 23°C. Cells were pelleted, suspended in 0.1 ml of 1xSDS loading buffer (60 mM Tris-Cl pH 6.8, 2% SDS, 5% glycerol, 100 mM β-ME) and lysed by heating at 95^°^C for 5 min. Cell debris was removed by centrifugation at 14,000 rpm in a microfuge at 23^°^C for 5 min. Yeast protein extracts were separated on protein gels containing 10% polyacrylamide and transferred to an Immobilon-P membrane (Millipore). The membranes were probed with anti-Myc antibodies (9E10, 1 µg/ml, Millipore) overnight in Blotto (10 mM Tris-Cl pH 7.5, 150 mM NaCl, 0.1% Tween-20, 5% dry milk) at 4^°^C and proteins were visualized by chemiluminescence (SuperSignal, Pierce).

### Protein binding assays

Recombinant His_6_-Cdh1 containing beads and MBP-Hsl1 were produced and purified from baculovirus-infected cells and *E. coli*, respectively, as described previously (Ostapenko *et al*., 2008). HA-Cdh1 was prepared by translation *in vitro* using the TNT^®^ T7 quick coupled transcription/translation system (Promega) according to the manufacturer’s instructions. In all His_6_-Cdh1-binding reactions, approximately 4.0 µg of His_6-_Cdh1 was on the beads. For HA-Cdh1 binding, 15 µl of the *in vitro* translated reaction was incubated with His_6_-Cdh1-beads and bound HA-Cdh1 was detected by immunoblotting with 12CA5 antibodies (1 µg/ml, Roche). Competition experiments between MBP-Hsl1 and HA-Cdh1 were conducted with 0.2 -1.0 µg MBP-Hsl1 pre-incubated with His_6_-Cdh1 beads for 15 minutes prior to the addition of the HA-Cdh1 from an *in vitro* translation reaction and a subsequent 30-minute incubation. Bound proteins were detected by immunoblotting with 12CA5 (1 µg/ml, Roche) and anti-MBP antibodies (0.5 µg/ml, New England Biolabs).

**Supplementary Table 1.**
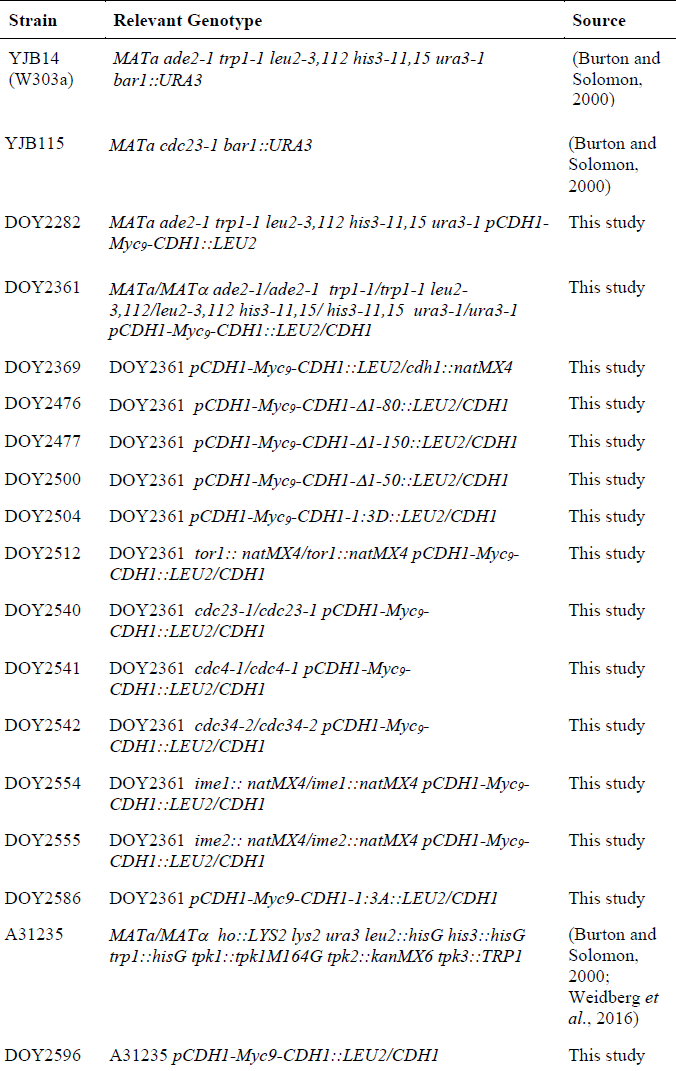
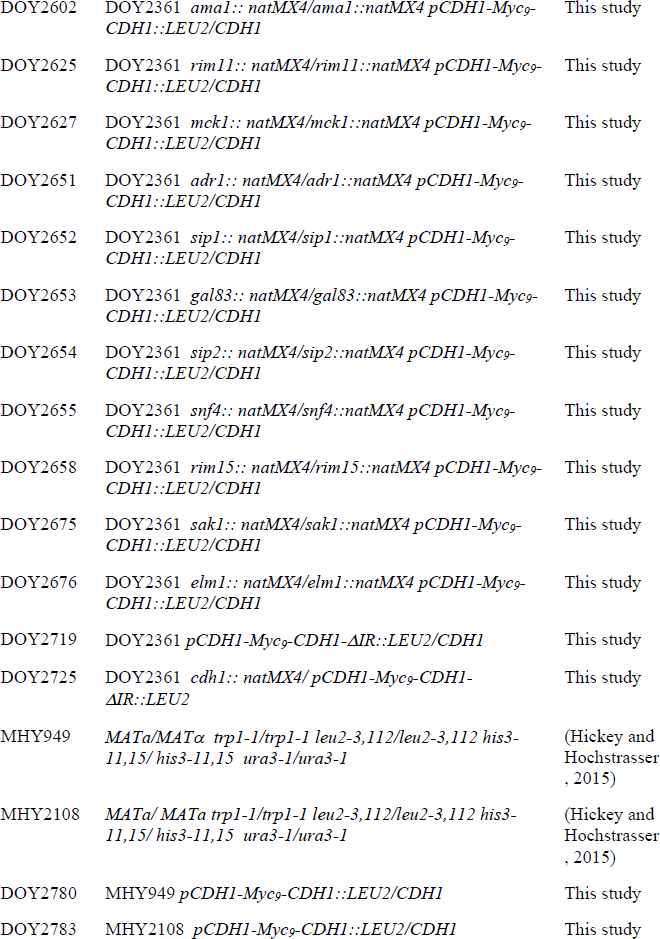
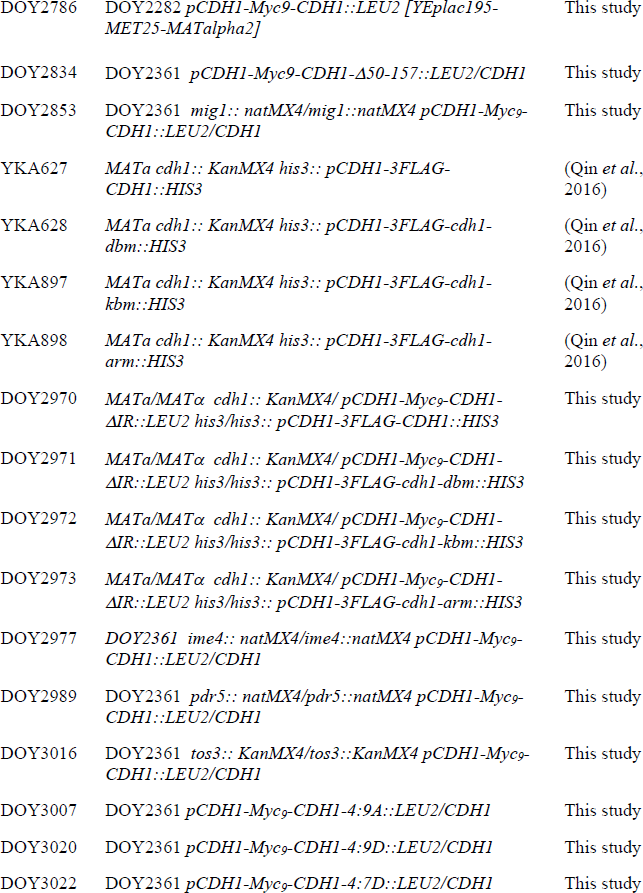
*S. cerevisiae* strains used in this study.

| Strain | Relevant Genotype | Source |
| --- | --- | --- |
| YJB14<br>(W303a) | *MATa ade2-1 trp1-1 leu2-3,112 his3-11,15 ura3-1 bar1::URA3* | (Burton and Solomon, 2000) |
| YJB115 | *MATa cdc23-1 bar1::URA3* | (Burton and Solomon, 2000) |
| DOY2282 | *MATa ade2-1 trp1-1 leu2-3,112 his3-11,15 ura3-1 pCDH1-Myc9-CDH1::LEU2* | This study |
| DOY2361 | *MATa/MATα ade2-1/ade2-1 trp1-1/trp1-1 leu2-3,112/leu2-3,112 his3-11,15/ his3-11,15 ura3-1/ura3-1 pCDH1-Myc9-CDH1::LEU2/CDH1* | This study |
| DOY2369 | DOY2361 *pCDH1-Myc9-CDH1::LEU2/cdh1::natMX4* | This study |
| DOY2476 | DOY2361 *pCDH1-Myc9-CDH1-Δ1-80::LEU2/CDH1* | This study |
| DOY2477 | DOY2361 *pCDH1-Myc9-CDH1-Δ1-150::LEU2/CDH1* | This study |
| DOY2500 | DOY2361 *pCDH1-Myc9-CDH1-Δ1-50::LEU2/CDH1* | This study |
| DOY2504 | DOY2361 *pCDH1-Myc9-CDH1-1:3D::LEU2/CDH1* | This study |
| DOY2512 | DOY2361 *tor1:: natMX4/tor1::natMX4 pCDH1-Myc9-CDH1::LEU2/CDH1* | This study |
| DOY2540 | DOY2361 *cdc23-1/cdc23-1 pCDH1-Myc9-CDH1::LEU2/CDH1* | This study |
| DOY2541 | DOY2361 *cdc4-1/cdc4-1 pCDH1-Myc9-CDH1::LEU2/CDH1* | This study |
| DOY2542 | DOY2361 *cdc34-2/cdc34-2 pCDH1-Myc9-CDH1::LEU2/CDH1* | This study |
| DOY2554 | DOY2361 *ime1:: natMX4/ime1::natMX4 pCDH1-Myc9-CDH1::LEU2/CDH1* | This study |
| DOY2555 | DOY2361 *ime2:: natMX4/ime2::natMX4 pCDH1-Myc9-CDH1::LEU2/CDH1* | This study |
| DOY2586 | DOY2361 *pCDH1-Myc9-CDH1-1:3A::LEU2/CDH1* | This study |
| A31235 | *MATa/MATα ho::LYS2 lys2 ura3 leu2::hisG his3::hisG trp1::hisG tpk1::tpk1M164G tpk2::kanMX6 tpk3::TRP1* | (Burton and Solomon, 2000; Weidberg *et al.*, 2016) |
| DOY2596 | A31235 *pCDH1-Myc9-CDH1::LEU2/CDH1* | This study |
| DOY2602 | DOY2361 *ama1:: natMX4/ama1::natMX4 pCDH1-Myc9-CDH1::LEU2/CDH1* | This study |
| DOY2625 | DOY2361 *rim11:: natMX4/rim11::natMX4 pCDH1-Myc9-CDH1::LEU2/CDH1* | This study |
| DOY2627 | DOY2361 *mck1:: natMX4/mck1::natMX4 pCDH1-Myc9-CDH1::LEU2/CDH1* | This study |
| DOY2651 | DOY2361 *adr1:: natMX4/adr1::natMX4 pCDH1-Myc9-CDH1::LEU2/CDH1* | This study |
| DOY2652 | DOY2361 *sip1:: natMX4/sip1::natMX4 pCDH1-Myc9-CDH1::LEU2/CDH1* | This study |
| DOY2653 | DOY2361 *gal83:: natMX4/gal83::natMX4 pCDH1-Myc9-CDH1::LEU2/CDH1* | This study |
| DOY2654 | DOY2361 *sip2:: natMX4/sip2::natMX4 pCDH1-Myc9-CDH1::LEU2/CDH1* | This study |
| DOY2655 | DOY2361 *snf4:: natMX4/snf4::natMX4 pCDH1-Myc9-CDH1::LEU2/CDH1* | This study |
| DOY2658 | DOY2361 *rim15:: natMX4/rim15::natMX4 pCDH1-Myc9-CDH1::LEU2/CDH1* | This study |
| DOY2675 | DOY2361 *sak1:: natMX4/sak1::natMX4 pCDH1-Myc9-CDH1::LEU2/CDH1* | This study |
| DOY2676 | DOY2361 *elm1:: natMX4/elm1::natMX4 pCDH1-Myc9-CDH1::LEU2/CDH1* | This study |
| DOY2719 | DOY2361 *pCDH1-Myc9-CDH1-ΔIR::LEU2/CDH1* | This study |
| DOY2725 | DOY2361 *cdh1:: natMX4/ pCDH1-Myc9-CDH1-ΔIR::LEU2* | This study |
| MHY949 | *MATa/MATα trp1-1/trp1-1 leu2-3,112/leu2-3,112 his3-11,15/ his3-11,15 ura3-1/ura3-1* | (Hickey and Hochstrasser , 2015) |
| MHY2108 | *MATa/ MATa trp1-1/trp1-1 leu2-3,112/leu2-3,112 his3-11,15/ his3-11,15 ura3-1/ura3-1* | (Hickey and Hochstrasser , 2015) |
| DOY2780 | MHY949 *pCDH1-Myc9-CDH1::LEU2/CDH1* | This study |
| DOY2783 | MHY2108 *pCDH1-Myc9-CDH1::LEU2/CDH1* | This study |
| DOY2786 | DOY2282 *pCDH1-Myc9-CDH1::LEU2 [YEplac195-MET25-MATalpha2]* | This study |
| DOY2834 | DOY2361 *pCDH1-Myc9-CDH1-Δ50-157::LEU2/CDH1* | This study |
| DOY2853 | DOY2361 *mig1:: natMX4/mig1::natMX4 pCDH1-Myc9-CDH1::LEU2/CDH1* | This study |
| YKA627 | *MATa cdh1:: KanMX4 his3:: pCDH1-3FLAG-CDH1::HIS3* | (Qin *et al.*, 2016) |
| YKA628 | *MATa cdh1:: KanMX4 his3:: pCDH1-3FLAG-cdh1-dbm::HIS3* | (Qin *et al.*, 2016) |
| YKA897 | *MATa cdh1:: KanMX4 his3:: pCDH1-3FLAG-cdh1-kbm::HIS3* | (Qin *et al.*, 2016) |
| YKA898 | *MATa cdh1:: KanMX4 his3:: pCDH1-3FLAG-cdh1-arm::HIS3* | (Qin *et al.*, 2016) |
| DOY2970 | *MATa/MATα cdh1:: KanMX4/pCDH1-Myc9-CDH1-ΔIR::LEU2 his3/his3:: pCDH1-3FLAG-CDH1::HIS3* | This study |
| DOY2971 | *MATa/MATα cdh1:: KanMX4/pCDH1-Myc9-CDH1-ΔIR::LEU2 his3/his3:: pCDH1-3FLAG-cdh1-dbm::HIS3* | This study |
| DOY2972 | *MATa/MATα cdh1:: KanMX4/pCDH1-Myc9-CDH1-ΔIR::LEU2 his3/his3:: pCDH1-3FLAG-cdh1-kbm::HIS3* | This study |
| DOY2973 | *MATa/MATα cdh1:: KanMX4/pCDH1-Myc9-CDH1-ΔIR::LEU2 his3/his3:: pCDH1-3FLAG-cdh1-arm::HIS3* | This study |
| DOY2977 | DOY2361 *ime4:: natMX4/ime4::natMX4 pCDH1-Myc9-CDH1::LEU2/CDH1* | This study |
| DOY2989 | DOY2361 *pdr5:: natMX4/pdr5::natMX4 pCDH1-Myc9-CDH1::LEU2/CDH1* | This study |
| DOY3016 | DOY2361 *tos3:: KanMX4/tos3::KanMX4 pCDH1-Myc9-CDH1::LEU2/CDH1* | This study |
| DOY3007 | DOY2361 *pCDH1-Myc9-CDH1-4:9A::LEU2/CDH1* | This study |
| DOY3020 | DOY2361 *pCDH1-Myc9-CDH1-4:9D::LEU2/CDH1* | This study |
| DOY3022 | DOY2361 *pCDH1-Myc9-CDH1-4:7D::LEU2/CDH1* | This study |

**Figure 4S.**
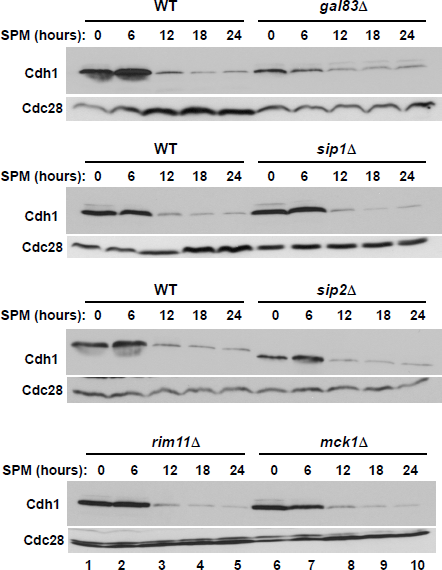
Roles of individual Snf1 pathway components in promoting Cdh1 turnover. Wild-type (strain DOY2361) and isogenic diploid yeast strains carrying deletions of individual SNF1 complex β-subunits, *GAL83* (strain DOY2653), *SIP1* (strain DOY2652) and *SIP2* (strain DOY2654); and meiosis-specific kinases *RIM11* (strain DOY2625) and *MCK1* (strain DOY2627), were grown in YPD and transferred to sporulation-inducing medium as in Figure 1A. Samples were withdrawn at the indicated times and probed to measure Cdh1 protein levels with 9E10 antibodies. The membranes were then re-probed with anti-PSTAIR antibodies to detect Cdc28 as a loading control.

